# Empirical methods for the validation of Time-To-Event mathematical models taking into account uncertainty and variability: Application to EGFR+ Lung Adenocarcinoma

**DOI:** 10.1101/2022.09.08.507079

**Authors:** Evgueni Jacob, Angélique Perrillat-Mercerot, Jean-Louis Palgen, Adèle L’Hostis, Nicoletta Ceres, Jean-Pierre Boissel, Jim Bosley, Claudio Monteiro, Riad Kahoul

## Abstract

**Background:** Over the past several decades, metrics have been defined to assess the quality of various types of models and to compare their performance depending on their capacity to explain the variance found in real-life data. However, available validation methods are mostly designed for statistical regressions rather than for mechanistic models. To our knowledge, in the latter case, there are no consensus standards, for instance for the validation of predictions against real-world data given the variability and uncertainty of the data. In this work, we focus on the prediction of time-to-event curves using as an application example a mechanistic model of non-small cell lung cancer. We designed four empirical methods to assess both model performance and reliability of predictions: two methods based on bootstrapped versions of parametric statistical tests: log-rank and combined weighted log-ranks (MaxCombo); and two methods based on bootstrapped prediction intervals, referred to here as raw coverage and the juncture metric. We also introduced the notion of observation time uncertainty to take into consideration the real life delay between the moment when an event happens, and the moment when it is observed and reported.

**Results:** We highlight the advantages and disadvantages of these methods according to their application context. We have shown that the context of use of the model has an impact on the model validation process. Thanks to the use of several validation metrics we have highlighted the limit of the model to predict the evolution of the disease in the whole population of mutations at the same time, and that it was more efficient with specific predictions in the target mutation populations. The choice and use of a single metric could have led to an erroneous validation of the model and its context of use.

**Conclusions:** With this work, we stress the importance of making judicious choices for a metric, and how using a combination of metrics could be more relevant, with the objective of validating a given model and its predictions within a specific context of use. We also show how the reliability of the results depends both on the metric and on the statistical comparisons, and that the conditions of application and the type of available information need to be taken into account to choose the best validation strategy.

## Introduction

Mechanistic models and by extension knowledge-based models provide a mathematical representation of biological phenomena, and by extension physiological and pathophysiological mechanisms. Based upon knowledge in the literature describing components of biology which are integrated using fundamental laws of nature such as physical and biochemical principles, these models allow representation and analysis of complex dynamic behavior of variables seen in biology and clinical trials ^1, 2^. During the past decade, mechanistic models have been progressively integrated into the pharmaceutical research and development industry workflow to provide valuable decision support in addition to conventional *in vitro* and *in vivo* approaches ^3, 4^.

An essential benefit of mechanistic models, when compared to statistical models or machine learning approaches, is that the model equations and associated parameters have a direct physical or biological meaning. Indeed, statistical models are based on the correlation found between variables while mechanistic ones model causality. This facilitates the overall comprehension of the process and the scientific interpretation of model results ^5^. Moreover, mechanistic modeling can predict biological or physical behaviors that have not yet been reported by currently available *in vivo* or *in vitro* experiments ^6 7^.

However, because of their complexity, and because this approach is more driven by knowledge, which can be considered as consolidated data, and less so by analysis of a small number of raw data from a very limited trial dataset, their credibility is often questioned compared to historical approaches, particularly their capacity to fully reproduce real-world data ^8 9^. For this reason, while the adoption of mechanistic modeling is in use at most major pharmaceutical/biotechnological companies, and its application is accelerating, trust in the relevance of such approaches for predicting novel phenomena is still a work-in-progress ^10 11 12^ but much work has been done and these models are currently used for decision-analytic (“GO/NOGO”) and process improvement (e.g. trial optimization) purposes. For example, models have been used to motivate the change of an “Approvable” to an “Approved” FDA decision for an oral anti-infective. They also have been used to successfully show, quantitatively and correctly, that the mechanism that a novel asthma therapy candidate was based upon was incorrect and that the therapy wouldn’t work. In another example, Pfizer found that a novel diabetes therapy would not reduce HbA1c sufficiently with respect to existing therapies. In the asthma and diabetes cases, the programs were terminated and in both cases competitors did a trial which aligned closely with the model predictions. The oral drug example was estimated to have saved $100million. The asthma and diabetes examples both saved about $30M trial costs, according to the company that did the modeling (and in the asthma case, according to the company that spent $30 million on the needless trial). This is why every major pharma company has a Quantitative Systems Pharmacology (QSP) group or at least QSP initiatives with service providers. We did not include these examples as there are many articles and cases in the QSP field.

Such mechanistic models can be of crucial importance ^13 14 15^ when it comes to helping and optimizing drug development. Indeed, because the models leverage large biological and medical knowledge for their structure and parameter values, only a limited amount of additional data is required to build an informed model that can be used to explore multiple settings (e.g. doses, regimens, patients characteristics) and choose the ones likely to work in clinical setting. As a consequence, it reduces (i) the ethical cost by limiting the number of patients exposed to a treatment setting that would not be efficient for them, as well as (ii) time and financial cost by limiting the number of trials and medical staff needed to gather preliminary data. Also, mechanistic models are adaptable to changes due to their modular nature: for instance, the addition of new candidate treatment to compare with golden standards of care.

For example, the model we present in this article focuses on EGFR-mutant lung adenocarcinoma and can be useful to create a synthetic control arm with a historical standard of care. This would allow to enroll real-world patients in the investigational arm, and thus maximize the benefit they can have from the treatment. Also, should a new treatment be added in the model, it would allow the in silico analysis of the patient subpopulations that would best benefit from the investigational treatment.

Even if the links between the variables of interest in these models are reported and justified in the literature, the range, the distribution and the correlation of their parameter values are difficult to evaluate. To overcome this problem, calibration is now a standard step in mechanistic model construction. Calibration can be defined as the search for a set of model parameter values that allows the model to reproduce a predefined set of behaviors and dynamics, observed in real life ^13^. However, how can we ensure that a model calibrated on several relevant datasets is good enough to be considered as validated and credible for its intended use?

Indeed, as with statistical models, mechanistic models have to be validated in order to confirm that their predictions are reliable and accurate. To avoid tautological bias and improve model credibility, this step requires data that has not been previously used for other purposes, such as model calibration ^16 17^. Model validation is a very topical issue, and is of interest to regulatory agencies. Indeed, the ASME V&V 40 Subcommittee on Verification and Validation (ASME V&V 40) in Computational Modeling of Medical Devices developed a risk-informed credibility assessment framework including a quantitative validation phase ^10^, and the European Medicines Agency (EMA) has drafted a specific guidance on the reporting of PBPK models including the evaluation of the predictive performance of the drug model ^18^. According to these guidelines, the context of use (CoU) of a model must also be defined, which defines the specific role and scope of the model in addressing the questions of interest ^19^.

The validation on retrospective data requires careful choice of appropriate metrics that take into account the nature of the measurements and the existing variability and bias ^20 21 22^. In the case of mechanistic models, the outputs are variable dynamics over time which are, most frequently, related to discrete reported observational or experimental values. Such experimental measures show an inherent variability due to the type of instruments that were used, as well as its resolution or sensitivity level, the quality of the sample, the applied protocol, human variability in reporting results and variability between samples ^23 24 25 26^, that also needs to be considered in the establishment of a validation strategy. In a situation of time-to-event (TTE) data, an additional difficulty can arise. Indeed, the TTE reported in real life corresponds to the moment when the event is detected by the observer, and not to the moment when it really happened. These two moments can be separated by a potentially significant period of time depending on the frequency of observations. Moreover, the model’s purpose is to predict the exact time until the occurrence of the event, and not the time to the observation of the event. This concept also has to be taken into consideration during the validation process.

Another goal of the validation process is also to guarantee that the model is not overfitted, which can happen if it was calibrated using datasets with limited variability. Additionally, the validation should as well assure that the variability of predictions of the model is not excessively wide. The latter can be assessed by evaluating the prediction intervals, that is to say, the range within which future observations should fall. Therefore, an adequate validation strategy should prevent both overfitting and underfitting as designing a model with the appropriate complexity requires achieving a balance between bias and variance, as well as a control of overfitting. Otherwise, if the prediction interval is too wide, the model’s outputs will lack precision and therefore the model’s credibility and usefulness will be low ^27^.

In order to demonstrate how the validation approach is applied, a case study including TTE oncological data will be described.

In summary, in this article we address the following challenge: how to properly manage the validation process when faced with a multi-condition situation, namely:

● Discrepancy in the size of the data to be compared: indeed, on the one hand, a mechanistic model may produce a very large amount of data. On the other hand, we wish to challenge the model outputs with a limited experimental validation dataset. Issues such as excess of statistical power, discrepancy in variability and uncertainty quantification will likely arise.
● Hypotheses for the application of statistical tests are not always verified producing a lack of statistical power.
● The uncertainty linked to the occurrence of events during clinical studies: the observation time uncertainty (further detailed in the “Methods” section), that is not handled by one deterministic model.

In this article, we focus on quantitative validation, a step of the overall validation process recommended by the regulatory guidelines (ASME V&V 40 ^28^ and EMA ^18^). We introduce multiple methods suited to validate deterministic non-linear mechanistic models including feedback loops, producing a TTE type of outcome. Importantly, these validation approaches consider both the model uncertainties and the variability of validation data.

We first present the methodology behind each one of those approaches, including the pre-processing of the dataset. Then, to answer the question of interest, we design four empirical methods to assess the model’s performance and the reliability of predictions: two methods based on bootstrapped versions of parametric statistical tests (log-rank and combined weighted log-ranks – MaxCombo) and two methods based on bootstrapped prediction intervals (that we named raw coverage and juncture metric). We also introduce the notion of observation time uncertainty (OTU) to take into consideration the delay between the moment when an event actually occurs, and the moment when it is witnessed and reported. Indeed in clinical routine, the time until an event of interest is not known precisely and instead, only is known to fall into a particular interval between two visits where the event could be reported by a clinician ^29 30^.

We then present an application on a clinical example. We finally discuss our results, highlight the advantages and disadvantages of these methods according to the application context, compare the performances and conclude.

## Methods

The statistical approaches, which are described hereafter, are combined with two additional mathematical concepts in order to better match the actual clinical context of this application example. Thus we first introduce the bootstrap and OTU concepts and then proceed with the actual statistical validation approaches.

### Bootstrap

In the context of modeling and simulation, one is not theoretically limited by the number of simulated statistical units (patients). This can be an advantage but also a drawback when using inferential statistics. Indeed, under the assumption of the same variance of the data, the statistical power will increase with the size of the sample ^31^. This can lead to a misinterpretation of the results if the test is statistically significant, concluding that there is a clinically relevant difference between compared groups when there is none ^32^. In order to control this statistical power, to avoid tests from being overly sensitive to negligible differences between groups, and to take into account the model uncertainty and the variance of the sampling in the simulation results, a bootstrapped version of the statistical tests is recommended^33 34 35^. By using a bootstrapped approach, the statistical tests are not applied to the entire population but to smaller samples (defined as being equal to the size of an ongoing clinical study (to mimic this study) or calculated a priori via the usual sample size calculation methods), therefore preventing the tests from being excessively sensitive because of an excess of statistical power ^36^. Repeating the calculations by bootstrapping makes it possible to empirically determine the distribution of the statistic which stabilizes around its value after a certain number of iterations according to the central limit theorem ^37^.

The output of the bootstrapped-testing approach is a ratio of significant or non-significant tests at a defined alpha risk (set to 5% in our case) out of the total number of performed tests ^38 39 40 41^. The ratio of significant tests in a context of bootstrapped testing can be linked to the empirical statistical power of the test ^42^. This ratio is then compared to a given threshold. For the sake of homogeneity with the two interval based approaches which will be introduced in the following sections, we will focus on the ratio of non-significant tests, meaning that the higher the ratio, the better the model predictions (no rejection of the null hypothesis). We chose to set the value of this threshold at 80%, as this value is conceptually the mirror of the currently accepted value for the power of a comparison when one wants to reject the null hypothesis ^43^ .

To determine how many iterations are required, preliminary tests are performed to see how long it takes for the ratio of non-significant tests to become stable (cf. supplementary file).

### Observation Time Uncertainty

The mechanistic models considered here are deterministic. Because we have access by design to the model outputs at all time points, the exact time at which a simulated event takes place can be determined. This model output is named the predicted-time-to-event (PTTE). In real patients, the true TTE can only be bounded between the time of two observations. We do not know the “exact” time-to-event. Therefore, an unknown difference between the PTTE and the reported time-to-event (RTTE) exists, bounded by the time between two observations (Figure 1). This time frame is what will be called the OTU, and depends on the delay between two observations. In other words, the actual TTE could have occurred in a time period ranging from the reported RTTE to the RTTE minus the time elapsed since the previous observation period. Nevertheless, one should keep in mind that we should not expect the model to cover this entire period since there is no evidence that the whole area reflects the real time at which the event occurred. A visual representation of the OTU is presented below.

**Figure 1:**
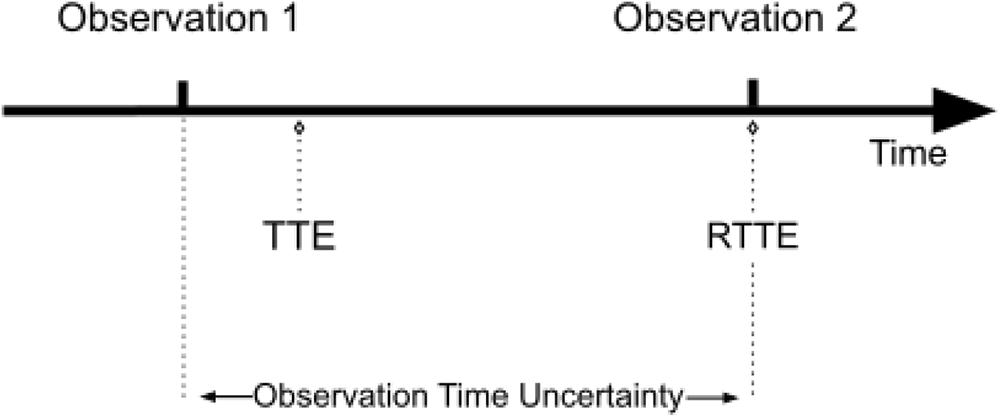
Representation of the observation time uncertainty. If an event happens between observation 1 and observation 2, it will only be reported at the time of observation 2. The observation time uncertainty corresponds to the time between two observations. To note, even though the TTE can theoretically happen any time between two observations, it is unknown whether this is true in a real life context. TTE: time-to-event, RTTE: reported time-to-event

The two concepts introduced above (bootstrap and observation time uncertainty) will be used in combination with the following validation approaches. Their description as well as their advantages and limits will be described.

#### Raw coverage

In order to perform both a quantitative and a visual validation of the computational model based on the validation dataset, a raw data coverage validation is performed. This approach consists of computing the percentage of the observed curve covered by the prediction interval of the model. In the context of simulation, there is no limit to the number of times one can run the same model, changing a few numbers of parameters and getting a new endpoint value. As a consequence, one can perform a large number of model runs so that there are more available model endpoints values than the number of endpoints values reported within the real population. Therefore, the definition of the prediction interval computed using bootstrapping has been adapted. At each iteration, a sample of simulated endpoints of the same size as the size of the real life population is taken from the set of simulated endpoints, and a Kaplan-Meier (KM) time to progression curve is estimated. This step is repeated *n* times. For each of these samples, based on the estimated Kaplan-Meier curve, an interpolation of the survival probability is then performed, for each of the times when an event is recorded in the entire simulated population. The distribution of the probability of being event-free is then computed for each time point, and based on the data collected from all samples, the empirical 2.5% and 97.5% quantiles are then calculated, the latter will serve as boundaries for the prediction interval. The level of coverage of the observed curve with the prediction interval of the simulated curve is then computed: for each time point, a check is performed to see if the observed curve is within the prediction interval – value is set to “True” – or not – value is set to “False” -. The percentage of “True” values is then computed. If the ratio is greater than a predetermined threshold, then, the model is considered to be validated. We estimated that a coverage value of at least 80%, meaning that no less than 80% of the observed survival curve is included in the prediction interval, is acceptable to consider the model as validated.

The choice of the right threshold is a critical process which involves biomodelers and biostatisticians, and has to answer certain criteria: (i) it has to be high enough to be restrictive, therefore demonstrating the model’s predictive capability, not excessively high so that (ii) it does not force the model to be overfitted on the validation dataset, (iii) it can be attained given the constraints of the validation process (e.g.: the quality of the available validation dataset), (vi) the threshold should be connected to well established statistical concepts when possible (with test-based approaches), and/or justified by clinical or biological considerations. In the case of interval based approaches such as the raw coverage, or the juncture metric which will be introduced in the next section, the validation threshold has been defined while having in mind that it can be difficult for a model to accurately predict a time-to-event variable at the very beginning and the very end of the observation period, also considering the fact that in those periods, model predictions have a usually narrow associated prediction interval, therefore justifying a threshold of 80% (the remaining 20% corresponding to the early and late periods of the observation window).

The raw coverage approach has the advantage of using the raw observed data without any prior transformation, considering that the real events occurred exactly at the moment of the reported event. In addition to the computed metric, the raw coverage allows one to easily perform a graphical check of the model’s ability to reproduce the observed results (see Figure 2 for an example of this approach applied to synthetic data).

The fact of considering that the event happened exactly at the time of the observation can also be considered as a limitation, as this is very unlikely. Indeed, the real event most certainly occurred sometime in between observation periods. Another point of concern is that the value of the raw coverage strongly relies on the width of the prediction interval. Indeed, the wider the interval, the more chances for the observed curve to be included in it. This means that if the model produces a lot of variability, then the raw coverage value will most certainly be very high.

**Figure 2:**
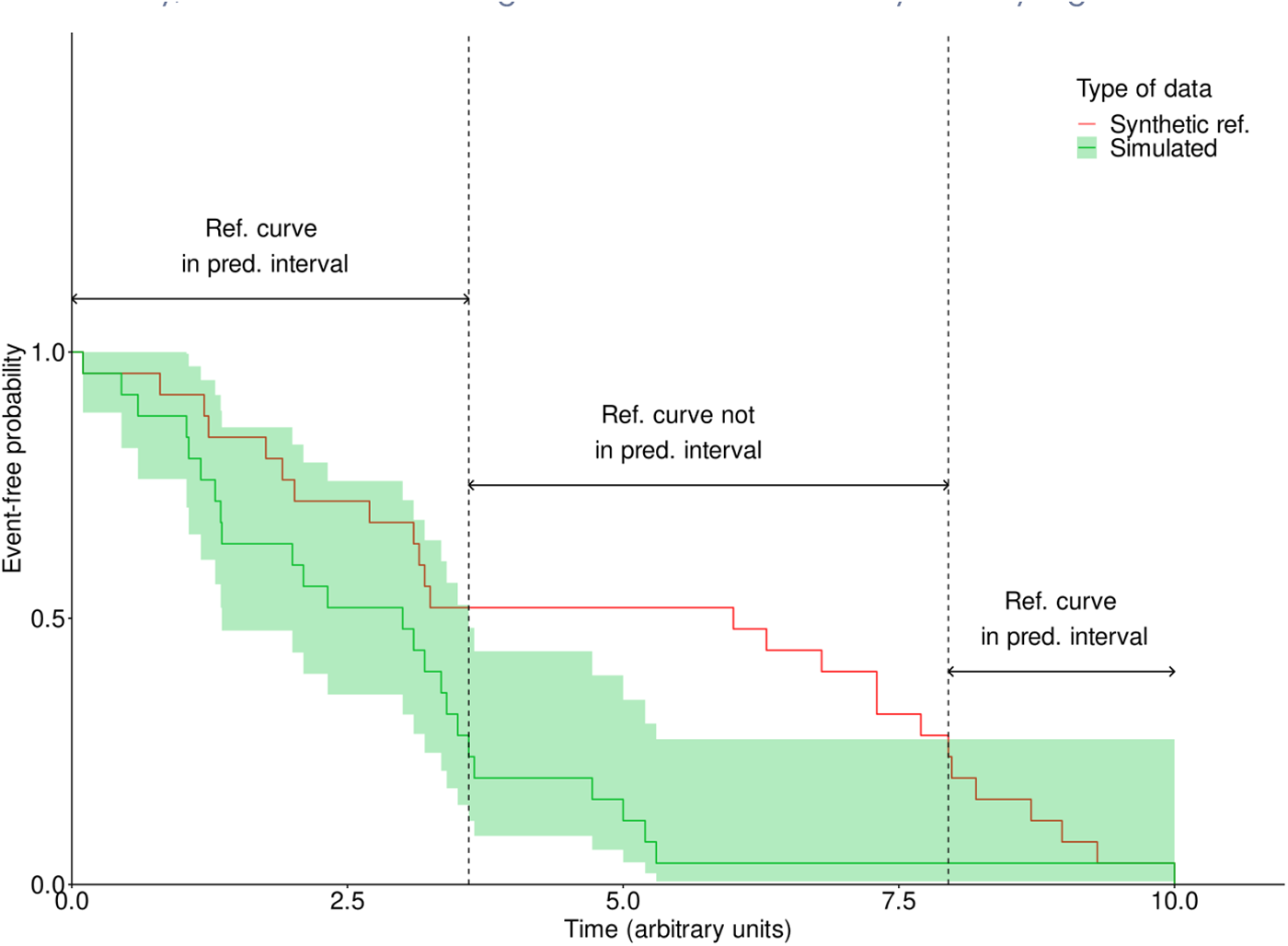
Representation of the raw coverage. The purpose of this example is to illustrate the raw coverage metric based on generated synthetic data. In the time interval of 0 to 10 months, the synthetic reference curve is covered by the prediction interval from t=0 to t=3.6 months, then between t=7.95 and t=10 months. This gives a raw coverage of ((3.6-0) + (10-7.95)) / (10-0) = 56.5%.

#### Juncture

The juncture approach is similar to the raw coverage in the way that it is both a mathematical and a visual validation method. It differs from the latter by the fact that it takes into account the OTU in the form of an interval and therefore does not rely on the assumption that the event occurred exactly at the time it was reported. This approach aims to evaluate if there is a reason to think that, for each given time point, the model simulations have no chance to get along with the reported data.

We name the observation variability interval the bound between the originally reported survival curve, and a curve where all events are shifted by a delay equivalent to one OTU. Thus each event factually occurring within this interval would be clinically reported on the corresponding survival curve.

The juncture approach measures the proportion of time over the entire observation period where the clinical evaluation interval and the 95% prediction interval overlap even if it is only partially. More precisely, at each time point where observed data is available, a check is performed to see if the two intervals contain common values. Should the condition be met, it means that the model is able to accurately explain the reported data based on the available information. Otherwise, it would mean that the model is unable to describe the reported data behavior.

The juncture approach metric corresponds to the ratio of time points where this condition is met, over the total number of time points. If the ratio is greater than a predetermined threshold, then, the model is considered to be validated (see Figure 3 based on generated synthetic data for illustrative purposes).

Similarly to the raw coverage, it is easy to identify the periods of time during the observation period where the simulation outputs successfully reproduce the observed data.

One of the limits of the metric associated with this approach is that it is strongly dependent on the width of both intervals, that is to say on the variability initially included in the computational model, which can come from the data used to calibrate it, as well as on the size of the OTU. Indeed, the larger the time in between observations, the wider the interval, and vice versa. Moreover, with the juncture approach, even a slight overlap of the two intervals is enough to be considered satisfactory, at a given time point. This means that overall, even if a small fraction of the observed data is covered by the simulated outputs over the entire observation period, then the entire prediction will be considered as validated, even if the latter is shifted up or down compared to the observations. As for the raw coverage, the juncture approach does not rely on a statistical test, with a p-value as an output, but instead on an arbitrary value between 0 and 100%. In a similar fashion, a value above 80% is considered to be acceptable to judge the model predictions as validated.

**Figure 3:**
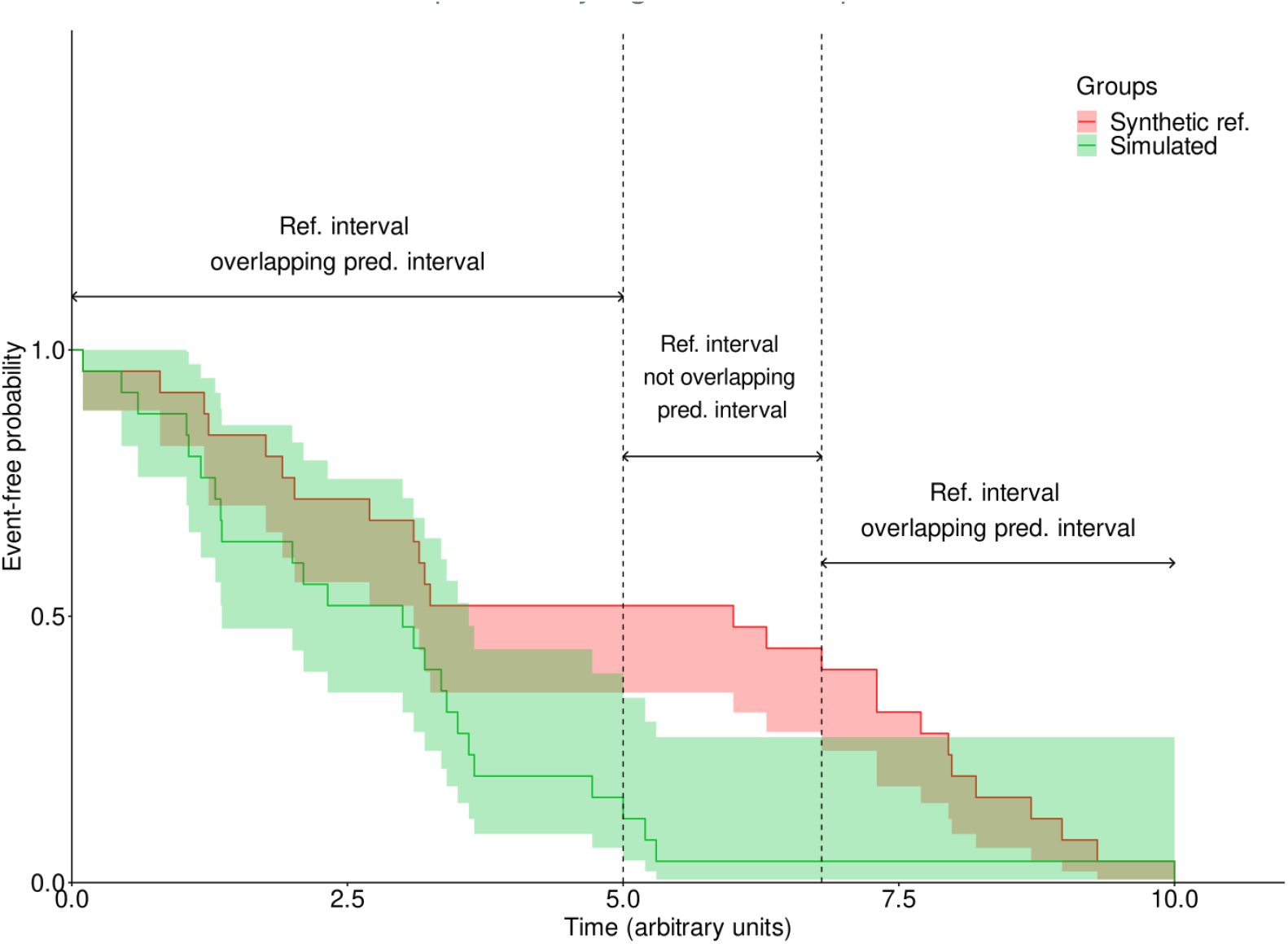
Representation of the juncture metric. The purpose of this example is to illustrate the “juncture” metric based on generated synthetic data. In the time interval of 0 to 10 months the synthetic reference interval overlaps, at least partially with the prediction interval from t=0 to t=5 months, then between 6.8 and 10 months. This results in a juncture of ((5-0) + (10-6.8)) / (10-0) = 82%.

#### Bootstrapped log-rank test

The log-rank is a well known and widely used test to compare survival curves ^41 44^. Its statistic is based on the computation of the difference between the observed and expected number of events in one of the groups at each observed event time. These differences are then added up to get an overall summary across all-time points where there is an event. The log-rank does not rely on the proportional hazards assumption, that is to say the risk associated with the event of interest remains proportional in both compared groups over the course of the follow-up period. It is a valid test of the null hypothesis of equality of survival functions. Nevertheless, if the proportional hazards assumption is not met, the log-rank test will be less powerful and therefore less capable to detect a difference between the two compared groups. ^45 46^.

Log-rank’s assumptions are the following: the degree of censoring should not be related to the outcome, and the events should have really happened at the reported time.

The log-rank test is integrated into a bootstrapped approach, and is tested first on the raw experimental data. It is then tested again with an OTU sampled from a uniform distribution *U*(-OTU, 0) being assigned to each real patient, at each iteration. A given number of bootstrap iterations are performed and the ratio of significant tests at a given alpha risk level is assessed. If this proportion does not exceed a certain predetermined threshold, the model is considered to be validated. As described in the “Bootstrap” section, a threshold of 80% has been chosen as the minimal value to be reached in order to consider the model’s predictions as validated with this method.

At each iteration proportional hazards assumption is checked, for exploratory purposes ^47 48^.

The advantage of the log-rank based validation approach is that it relies on a statistical test frequently used to assess differences between two samples when it comes to TTE data, making its results easy to understand. The proposition to combine the log-rank test with a bootstrap approach, with a sampling of a number of model runs comparable to the number of real patients, prevents it from being excessively sensitive to differences between groups because of an excess of statistical power induced by a very large number of statistical units.

However, because the statistical power of the log-rank test is affected by the proportional hazards assumption, its results might not be considered reliable for samples where the assumption is not met ^45^. This implies that if the number of samples where the proportional hazards assumption is not met is high, the ratio of significant tests can be biased. A method more suited for the situations where the proportional hazard hypothesis is not met is introduced in the next section.

In the case where the OTU is not taken into account, the TTE curve based on the sample taken from the simulated data is directly compared to the raw observed data, implying that the reported (RTTE) and the real TTE are equal, which can be considered as a strong assumption.

#### Bootstrapped weighted log-rank tests combination

Several statistical methods have been developed to better manage the risk of type 1 error and to optimize statistical power in a situation where the proportional hazard assumption is not met ^45 49 50 51^. One of these methods is the use of a combination of weighted log-rank tests, called the MaxCombo approach ^52 53^. This approach consists in the use of the Flemming-Harrington family of weights (FH(ρ, γ), ρ, γ ≥ 0). The combination of weights that is used is the following:

- FH(0,0) corresponding to a regular non-weighted log-rank
- FH(1,0) for a log-rank putting more weight on early differences
- FH(1,1) where weights are put on mid-observation differences
- FH(0,1) where late differences are given more weight

Similarly to the log-rank, this approach is also bootstrapped. At each iteration of the bootstrap, all four tests are performed, and the one with the highest z-score, that is to say, the test with the weights showing the largest difference between KM curves, is selected. Given the fact that four tests are performed at once, a Bonferroni correction is applied to the p-value of this test.

The weighted log-rank combination approach is a more robust version of the standard log-rank based one, usable even in the situation where the two compared survival curves cross, implying the proportional hazards assumption is not met. By drawing sub-samples, from the very large simulated population, of the same size as the observed population (a reasonable sample size), we control the occurrence of excessive statistical power related to large samples, preventing the test from being overly sensitive to neglectable differences between the two groups.

As out of the four tests performed at each iteration, it is the one with the highest z-score that is selected, this approach tends to find more differences than a standard log-rank because more weight is put on the time period where the distance between KM curves is at its maximum. For this reason, the validation acceptance threshold has to be defined accordingly, and should eventually be set lower when compared to other approaches. Similarly to the bootstrapped log-rank test, this approach is launched twice, first without the OTU, and a second time with a random OTU assigned to real patients at each iteration.

The advantages and limits of the 4 validation methods as well as their variants with OTU are summarized in the table 1 below.

**Table 1:**
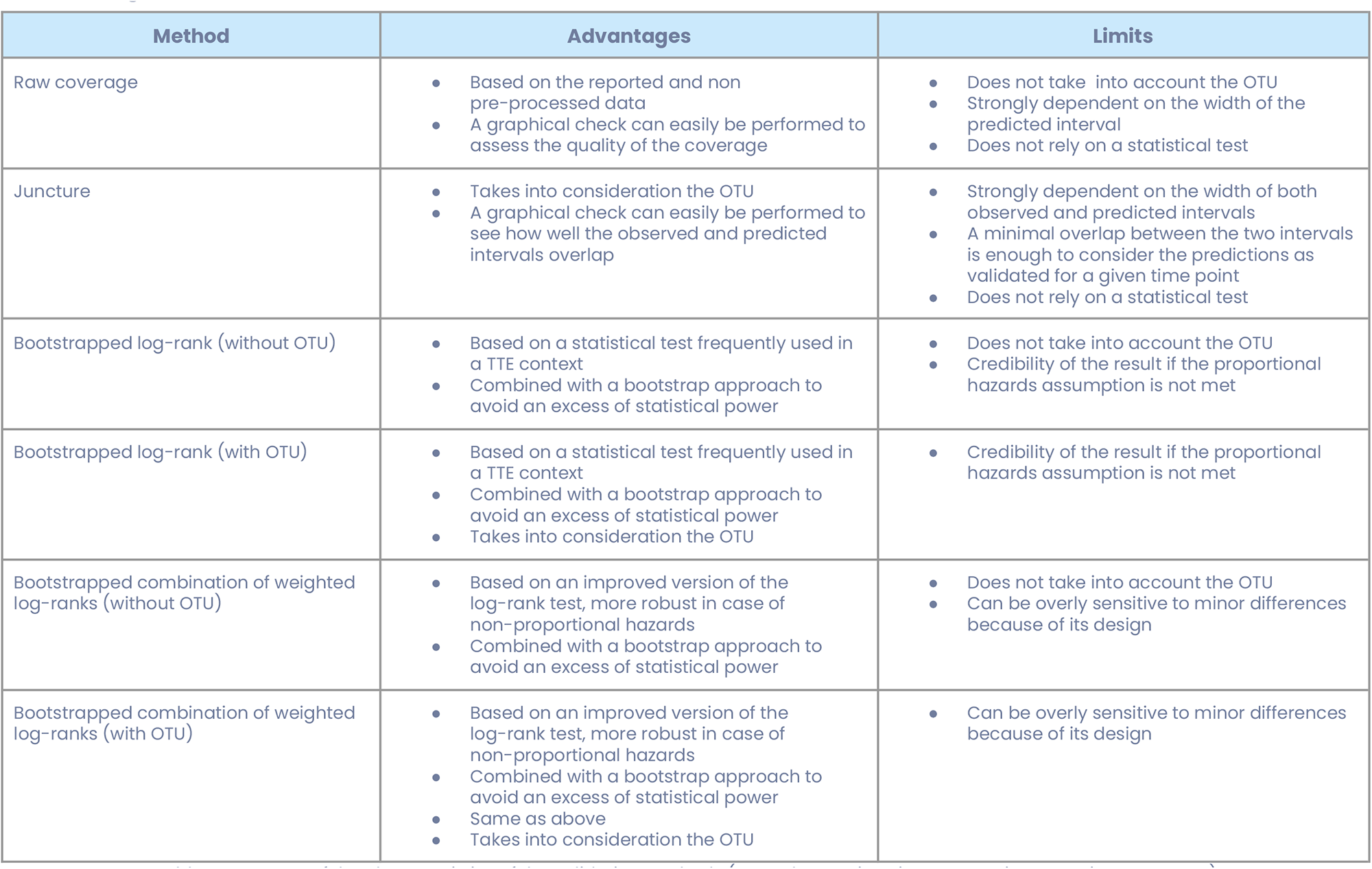
Summary of the characteristics of the validation methods (OTU: Observation Time Uncertainty, TTE: Time-To-Event)

| Method | Advantages | Limits |
| --- | --- | --- |
| Raw coverage | Based on the reported and non pre-processed data<br>A graphical check can easily be performed to assess the quality of the coverage | Does not take into account the OTU<br>Strongly dependent on the width of the predicted interval<br>Does not rely on a statistical test |
| Juncture | Takes into consideration the OTU<br>A graphical check can easily be performed to see how well the observed and predicted intervals overlap | Strongly dependent on the width of both observed and predicted intervals<br>A minimal overlap between the two intervals is enough to consider the predictions as validated for a given time point<br>Does not rely on a statistical test |
| Bootstrapped log-rank (without OTU) | Based on a statistical test frequently used in a TTE context<br>Combined with a bootstrap approach to avoid an excess of statistical power | Does not take into account the OTU<br>Credibility of the result if the proportional hazards assumption is not met |
| Bootstrapped log-rank (with OTU) | Based on a statistical test frequently used in a TTE context<br>Combined with a bootstrap approach to avoid an excess of statistical power<br>Takes into consideration the OTU | Credibility of the result if the proportional hazards assumption is not met |
| Bootstrapped combination of weighted log-ranks (without OTU) | Based on an improved version of the log-rank test, more robust in case of non-proportional hazards<br>Combined with a bootstrap approach to avoid an excess of statistical power | Does not take into account the OTU<br>Can be overly sensitive to minor differences because of its design |
| Bootstrapped combination of weighted log-ranks (with OTU) | Based on an improved version of the log-rank test, more robust in case of non-proportional hazards<br>Combined with a bootstrap approach to avoid an excess of statistical power<br>Same as above<br>Takes into consideration the OTU | Can be overly sensitive to minor differences because of its design |

### Application example: validation of a mechanistic model of lung adenocarcinoma under gefitinib treatment

The methods that were presented in the previous section were assessed and tested on a knowledge-based mechanistic model of the tumor evolution of patients with lung adenocarcinoma.

This model, named the *In Silico* Epidermal growth factor receptor Lung Adenocarcinoma (ISELA), evaluates tumor growth and progression in patients harboring a mutation on the Epidermal Growth Factor Receptor (EGFR), and relies on a mechanistic representation of the lung adenocarcinoma (LUAD) evolution from specific EGFR mutations to clinical outcome. It includes shrinkage in response to the administration of a first generation tyrosine kinase (TKI) drug called gefitinib. This model was calibrated with publicly available data ^54 55 56 57 58 59^, and details regarding the calibration of tumor growth are given in a paper published by Palgen *et al.* ^60^. It should be noted that this model is not designed to predict mortality from any cause, but rather developed to predict time to tumor progression (TTP), which was deduced from progression-free and overall survival curves.

In this application context, we focus on the TTP clinical endpoint and will apply our validation strategy to ensure the ISELA model’s accuracy on a dataset that was not previously used in the calibration process: the one extracted from Maemondo et *al.* and not previously used for calibration purposes ^61^. This study compares the effect of gefitinib versus chemotherapy on NSCLC (of which 90.4% are LUAD) with mutated EGFR. The trial described in the article, and called NEJ002, took place in Japan and gefitinib was used as the first-line treatment. About 90% of the analyzed population had stage IIIb or IV cancers. In this study, gefitinib (250 mg/d) was orally administered once daily, until disease progression, development of intolerable toxic effects, or withdrawal of consent. The progression-free survival (PFS) and the overall survival (OS) curves were manually extracted for patients treated with gefitinib.

#### Pre-processing of the datasets

A gap was identified between the model output and the dataset related endpoint. While the model represents TTP, which is a clinical endpoint that censors out the patients that die, the dataset extracted from Maemondo et *al.* focuses on PFS and OS. In both clinical endpoints, a patient’s death prior to disease progression is therefore an event and is not censored out.

To be able to compare the model TTP to the experimental dataset, the endpoint disease progression was derived from clinical PFS and OS: we manually extracted the KM curves of PFS and OS and their corresponding censored events, and deduced the list of PFS and OS TTEs.

Under the assumption that patients who died before disease progression are characterized by the same time to event in the PFS and OS sets, we are able to filter out PFS events that correspond to patients’ death. Indeed, by removing from the PFS values all TTEs that are equal in PFS and OS datasets with a small tolerance due to manual extraction uncertainty, one is left with the TTEs where events are disease progression only. The reduced dataset was named NEJ002 TTP. We consider as equal any PFS and OS values that differ from maximum 2 days.

The NEJ002 TTP dataset is composed of 74 patients, corresponding to 68% of the original dataset, a percentage which seems plausible, considering that the remaining 32% correspond to either censoring, or dead patients. Nevertheless, the exact number was not reported in Maemondo et *al.*. Among the removed data points, 24 correspond to censored events and 10 to death preceding disease progression. Removal of those data points leads to a shift of the curve towards the left. It should be noted nonetheless that the overall linear slope is unchanged (Figure 4).

The statistical validation methods described previously were applied to compare the ISELA simulation results to the NEJ002 TTP dataset. For all situations where a bootstrap approach was used, 5000 iterations were performed (cf. Appendix 1), while for approaches based on the log-rank test, the alpha risk level was set at 5%. The time between visits being 2 months, the OTU used ranged between -2 and 0 months. Note that the ISELA model represents the tumor growth from which we can deduce the TTP, and only right censoring can be represented by the model.

**Figure 4:**
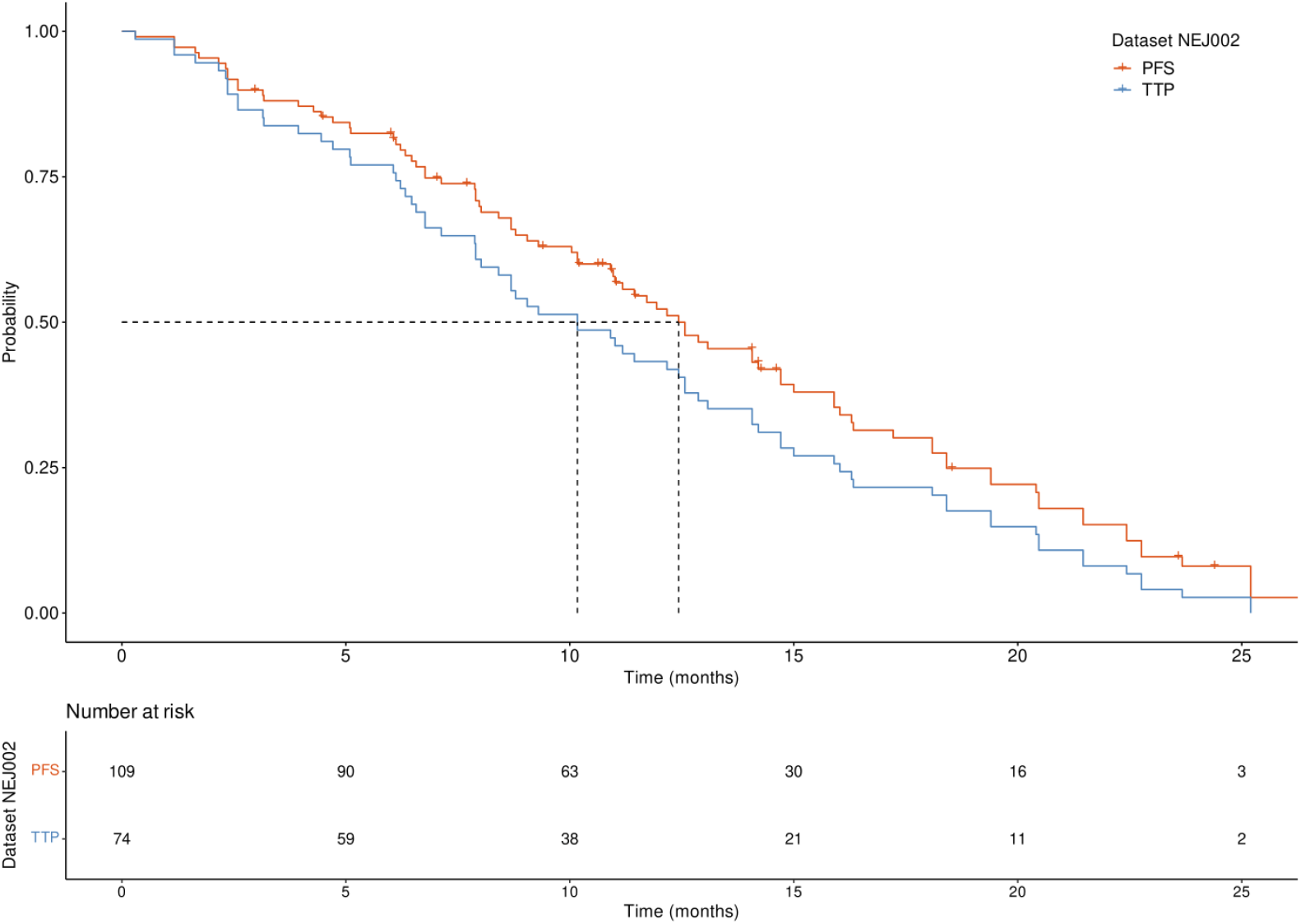
Survival curve based on the NEJ002 dataset. Probability of progression-free survival (red curve) and tumor non-progression (blue curve) respectively before and after removal of dead and censored patients. The dashed line highlights the impact of data-processing on time corresponding to the median probability. Median PFS (12.43 months) and TTP (10.17 months) are represented with dotted lines. PFS data manually extracted from Maemondo et al., processed and plotted in R.

## Results and discussion

### Results on entire population

According to the initially defined CoU, the validation approaches were applied to the data corresponding to the entire population extracted from the Maemendo et *al.* article and based on the NEJ002 trial. The results are shown in Fig. 5 and summarized in Table 2.

**Figure 5:**
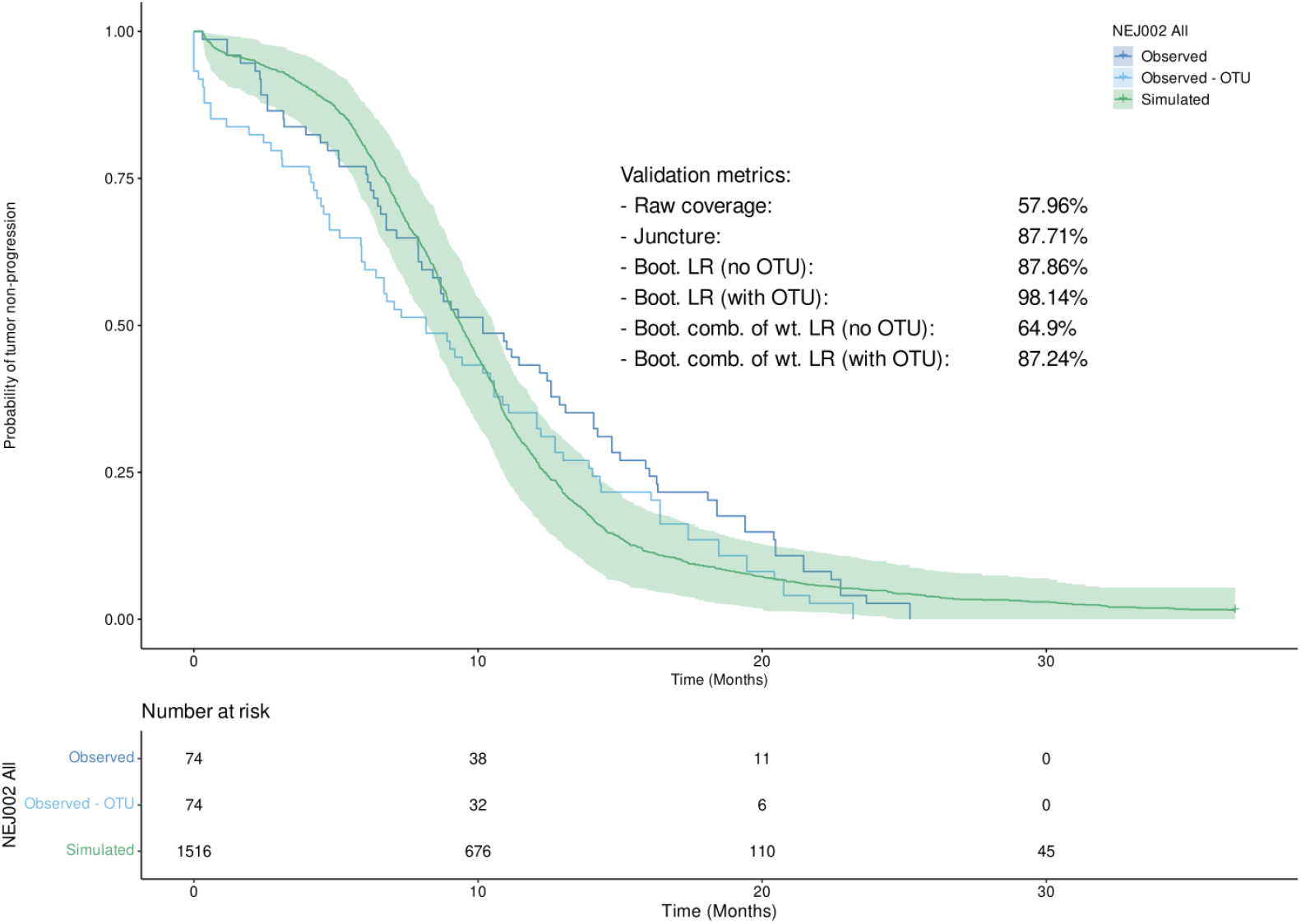
Observed and simulated Kaplan-Meier curves computed on the full dataset. The 95% bootstrapped prediction interval of the simulated curve is represented by the green area. (Boot. = Bootstrapped, LR = log-rank test, comb. of wt. LR = combination of weighted log-rank tests (MaxCombo))

**Table 2:** Results of the various validation methods applied to the full dataset. (PH = proportional hazards) Note: the acceptability threshold was set at 80%. Given the way all four metrics are defined, the higher the value, the closer the model predictions are to the observed values according to the validation assumptions.

| Method | Metric value | Ratio of of samples where PH assumption is not met |
| --- | --- | --- |
| Raw coverage | 57.96% | NA |
| Juncture | 87.71% | NA |
| Bootstrapped LR (no OTU) | 87.68% | 4.68% |
| Bootstrapped LR (with OTU) | 98.14% | 12.92% |
| Bootstrapped weighted log-rank combo (no OTU) | 64.9% | 4.62% |
| Bootstrapped weighted log-rank<br>combo (with OTU) | 87.24% | 13.28% |

In this context of use, the results provided by the various validation methods vary from 57.96% to 98.14% of validation. Four methods show a metric superior to the chosen threshold of acceptance set at 80%, while the two others fail to reach it. The raw coverage, and the weighted LR based method without OTU fail to reach the validation threshold. The reason for the raw coverage metric to remain below 80% can be explained by the fact that between 2 and 6 months, the model underestimates the number of events, and then overestimates them between 12 and approximately 24 months, as shown in Figure 5. Regarding the weighted LR based approach without OTU, it shows that the model’s predictions are not accurate while both bootstrapped LR metrics, with and without OTU, as well as the weighted LR with OTU, indicate that the model is performing well. This difference can be explained by the fact that simulated and observed curves cross, implying that the statistical power of LR tests is reduced, resulting in a lower rejection rate of the null hypothesis, and consequently, a higher validation metric. The fact that the OTU is taken into consideration also has a positive impact.

### Refinement of the context of use

According to previous results and the noticeable discrepancies between methods, in order to show how data structure and the model’s CoU can have an impact on the model validation process we decided to go further through the exploration of the data. Indeed, considering the mutational status of the tumor, the data used for validation consist of a mixture of two populations. Each of these subsets was characterized by a specific EGFR mutation: exon 19 deletion (Del19) and L858R on exon 21. Those mutations had an impact on the time to progression ^62 63 64^, making the simultaneous validation on both types of patients not relevant and potentially incorrect. Thus, in order to have a more precise assessment of the model’s predictive capability, the validation process assessment was stratified according to the mutation status of patients. This was done consistently with the calibration which was performed on a set of individual data patients, for which the EGFR mutation was specified.

After applying the validation approaches to the Del19 subset, new metrics were computed and summarized in Figure 6 and Table 3.

**Figure 6:**
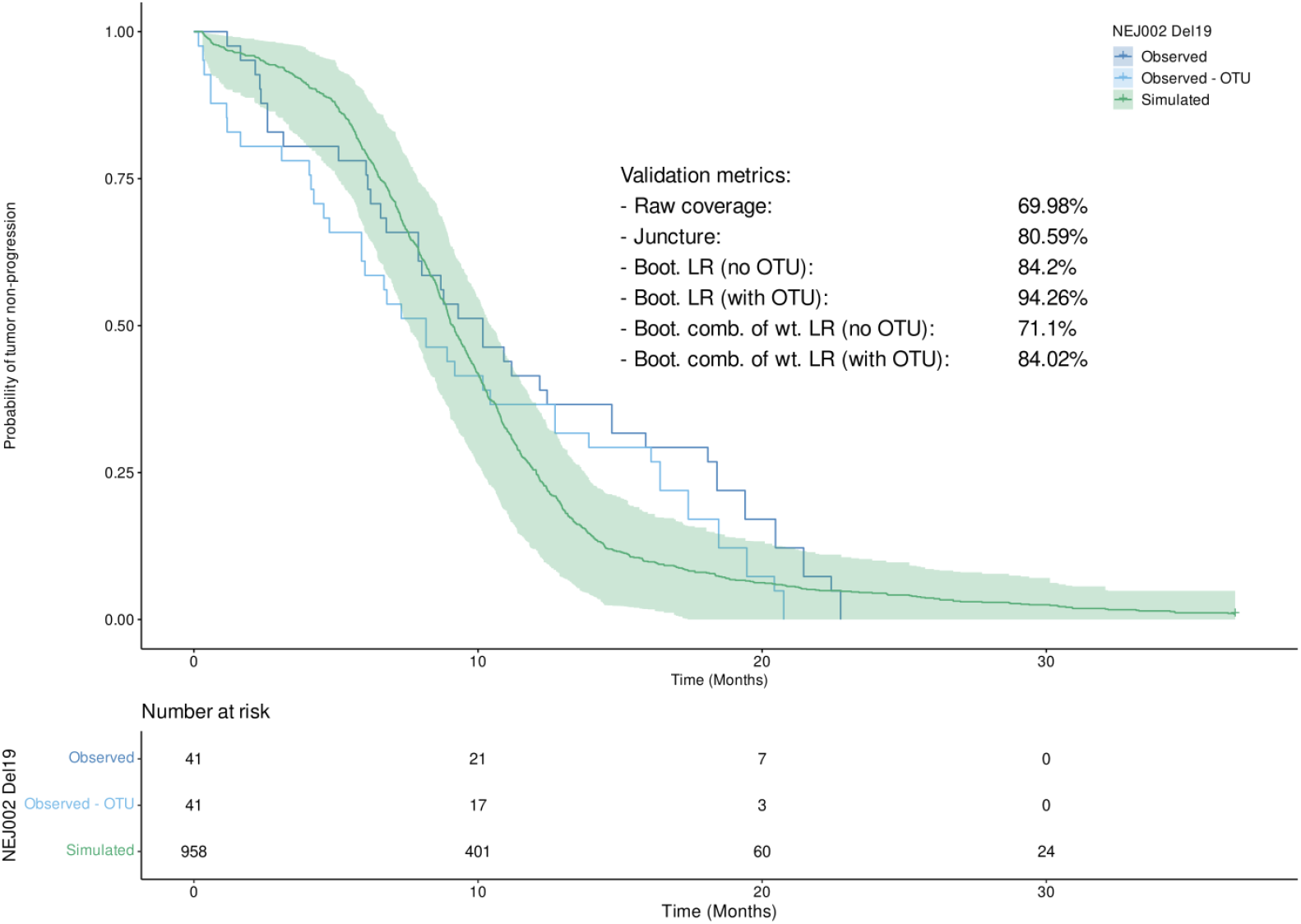
Observed and simulated Kaplan-Meier curves computed on the Del19 subpopulation. The 95% bootstrapped prediction interval of the simulated curve is represented by the green area. (Boot. = Bootstrapped, LR = log-rank test, comb. of wt. LR = combination of weighted log-rank tests (MaxCombo))

**Table 3:** results of the various validation methods applied to the Del19 subset. (PH = proportional hazards)

| Method | Metric value | Ratio of of samples where PH assumption is not met |
| --- | --- | --- |
| Raw coverage | 69.98% | NA |
| Juncture | 80.59% | NA |
| Bootstrapped LR (no OTU) | 84.2% | 12.34% |
| Bootstrapped LR (with OTU) | 94.26% | 19.24% |
| Bootstrapped weighted log-rank combo (no OTU) | 71.1% | 12.42% |
| Bootstrapped weighted log-rank combo (with OTU) | 84.02% | 19.02% |

As for the Del19 subset, the validation metrics on the L858R subset were computed and summarized in Figure 7 and Table 4.

**Figure 7:**
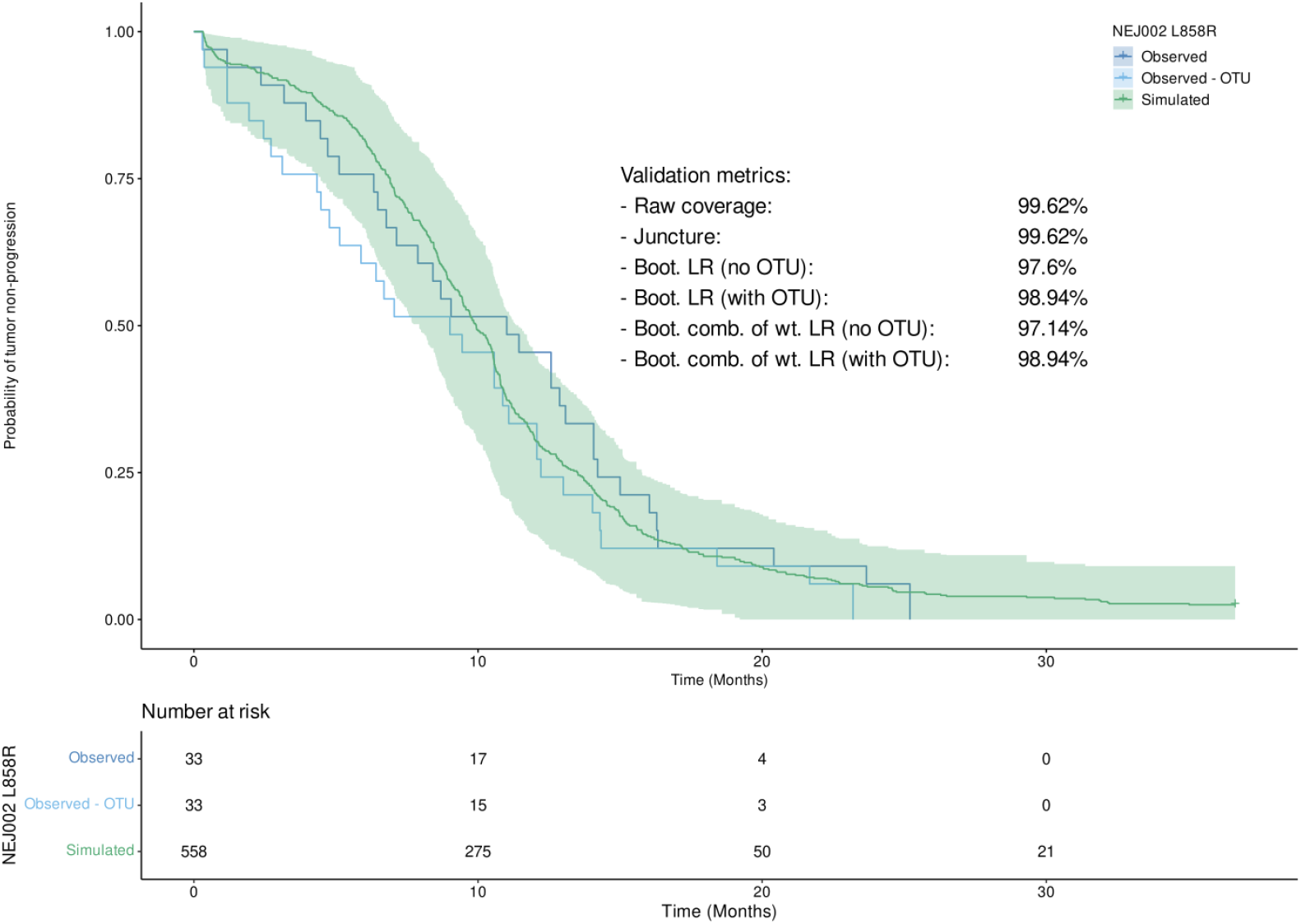
Observed and simulated Kaplan-Meier curves computed on the L858R subset. The 95% bootstrapped prediction interval of the simulated curve is represented by the green area. (Boot. = Bootstrapped, LR = log-rank test, comb. of wt. LR = combination of weighted log-rank tests (MaxCombo))

**Table 4:**
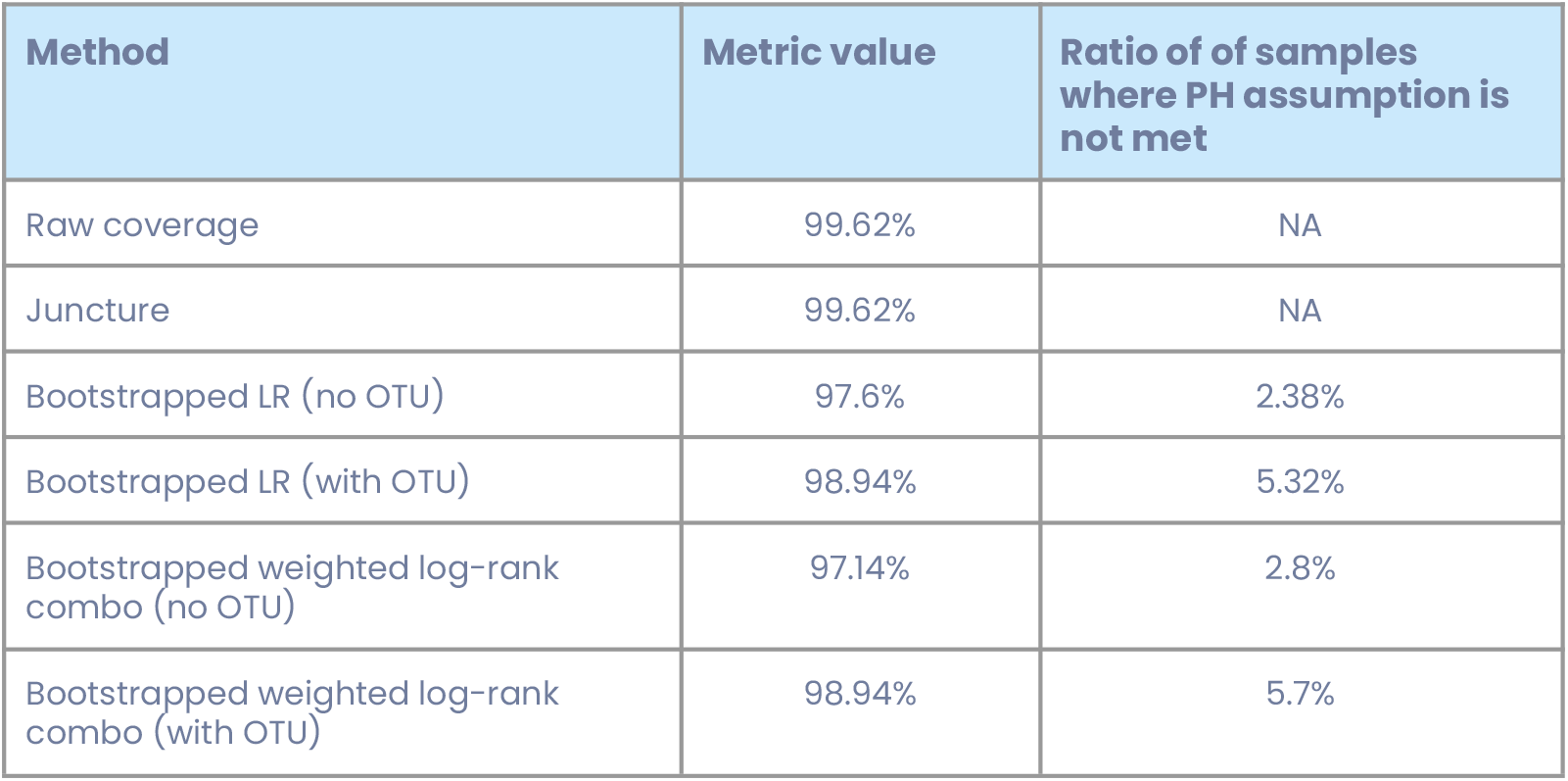
Results of the various validation methods applied to the L858R subset. (PH = proportional hazards)

When applied to the Del19 subset, the raw coverage approach provided better results than on the overall population with approximately 12% more coverage of the observed curve. Regarding the juncture method, the value was lower by 7.12% for the subset. A decrease was found as well for both the bootstrapped log-rank (-3.66% without OTU, -3.88% with OTU) and the bootstrapped combination of weighted log-ranks with OTU (-3.22%). The version without the OTU increased by 6.2%. The results obtained on the Del19 subset show that there are even more differences between the validation data and the simulations than in the previous CoU. The model appears to be unable to correctly predict events in this subgroup, despite the better results obtained with the raw coverage approach, which indicate that relying on a single metric is not enough to properly evaluate the quality of the model’s predictions. We note here, as an aside, that mismatches such as this help guide model improvement, allowing us to better understand the disease and treatments effects. Without such a model, these discrepancies might not even be noticed.

In the case of the L858R subset, both the raw coverage and juncture methods produced much better results than on the entire population : 99.62% for both approaches, equal to an increase of 29.64% and 19.03%, respectively. With the bootstrapped log-rank, the results were better without the OTU (+13.4%), as well as with the OTU taken into account (+4.68%). For the bootstrapped combination of weighted log-ranks, both metrics without and with OTU were better on the subset than on the global population (+32.24% and +14.92%, respectively). This demonstrates that all validation metrics can show good performances when the CoU is properly chosen. Indeed, it appears that the model is well suited to predict the events in the L858R subgroup, which was not the case for the Del19 subset.

The differences between the metrics obtained on the entire population and on the subsets are summarized in the table 5 below.

**Table 5:** Differences between the results obtained on the initial dataset and the Del19 and L858R subsets.

| Method | Difference between full dataset and Del19 | Difference between full dataset and L858R | Difference between Del19 and L858R datasets |
| --- | --- | --- | --- |
| Raw coverage | +12.02% | +41.66% | +29.64% |
| Juncture | -7.12% | +11.91% | +19.03% |
| Bootstrapped LR (no OTU) | -3.66% | +9.74% | +13.4% |
| Bootstrapped LR (with OTU) | -3.88% | +0.8% | +4.68% |
| Bootstrapped weighted log-rank combo (no OTU) | +6.2% | +36.24% | +32.24% |
| Bootstrapped weighted log-rank combo (with OTU) | -3.22% | +11.7% | +14.92% |

## Conclusion

In this article, we introduced different approaches to validate mathematical model predictions on Time-To-Event data, and gave some insight on how to perform a robust validation of a mathematical model by choosing one or multiple methods to correctly evaluate the model’s prediction. We have emphasized that the choice of methods and metrics is highly impactful and thus it should be made according to the context, available validation data, and to its specificities, structure and nature (single curve or interval).

We demonstrated in the application section that a model is meant to be applied to a specific context of use (CoU), as otherwise, by performing a validation on an excessively broad dataset, the whole process may fail because the model will not be able to correctly predict the events for heterogeneous subpopulations. Moreover, the importance of using multiple validation methods at once instead of relying on a single one was illustrated by the results obtained on a non-adapted CoU (*e.g.* Del19) where, by looking at only one validation metric (raw coverage in this specific case), one could wrongfully conclude that the model performed well, or at least better than within the previous CoU, while in fact, all the other metrics together demonstrated a worse performance of the model.

Indeed, the strength of the validation process comes from the combination of well selected validation metrics, as each one has its own strengths and weaknesses and conditions of application (*see* Table 1). We highlighted and suggested that simultaneously using multiple methods that rely on different statistical concepts can ensure correct evaluation of the model’s performance. Nevertheless, we noted that some of the methods introduced in this article will have more weight than others in a combined approach, because some methods are relatively more robust and more prone to detect differences between observed and simulated data, for example the MaxCombo approach when the assumption of proportional hazard is not met.

It should be noted that during the validation process, it is necessary to avoid tautology, principally by using data not used for the construction of the model, and to avoid trying to validate the model by arbitrarily changing goals, but to define a priori protocol and methods in order to evaluate the model and the context of use properly.

## Declarations

### Ethics approval and consent to participate

Not applicable.

### Consent for publication

Not applicable.

### Availability of data and materials

The datasets used and/or analysed during the current study are available from the corresponding author on reasonable request.

### Authors’ contributions

- EJ: Conceptualization, Methodology, Software, Validation, Formal analysis, Writing - Original Draft, Writing - Review & Editing, Visualization, Supervision
- APM: Conceptualization, Methodology, Software, Validation, Formal analysis, Writing - Original Draft, Writing - Review & Editing, Visualization, Supervision
- JLP: Conceptualization, Validation, Formal analysis, Methodology, Investigation, Visualization, Writing - Review & Editing
- AL: Conceptualization, Data curation, Formal analysis, Methodology, Writing - Review & Editing, Project administration
- NC: Data curation, Software
- JPB: Writing - Review & Editing
- JB: Writing - Review & Editing
- CM: Conceptualization, Methodology, Review & Editing, Project administration
- RK: Conceptualization, Methodology, Validation, Writing - Original Draft, Writing - Review & Editing

All authors read and approved the final manuscript.

## Acknowledgements

Not applicable.

