## Supplementary file for "Empirical methods for the validation of Time-To-Event mathematical models taking into account uncertainty and variability: Application to EGFR+ Lung Adenocarcinoma"

**Supplementary Figures**

**Figure S1**

**Evolution of the ratio of non-significant bootstrapped log-rank tests on the entire population according to the number of bootstrap iterations.** The ratio can be considered as stable after 5000 iterations.

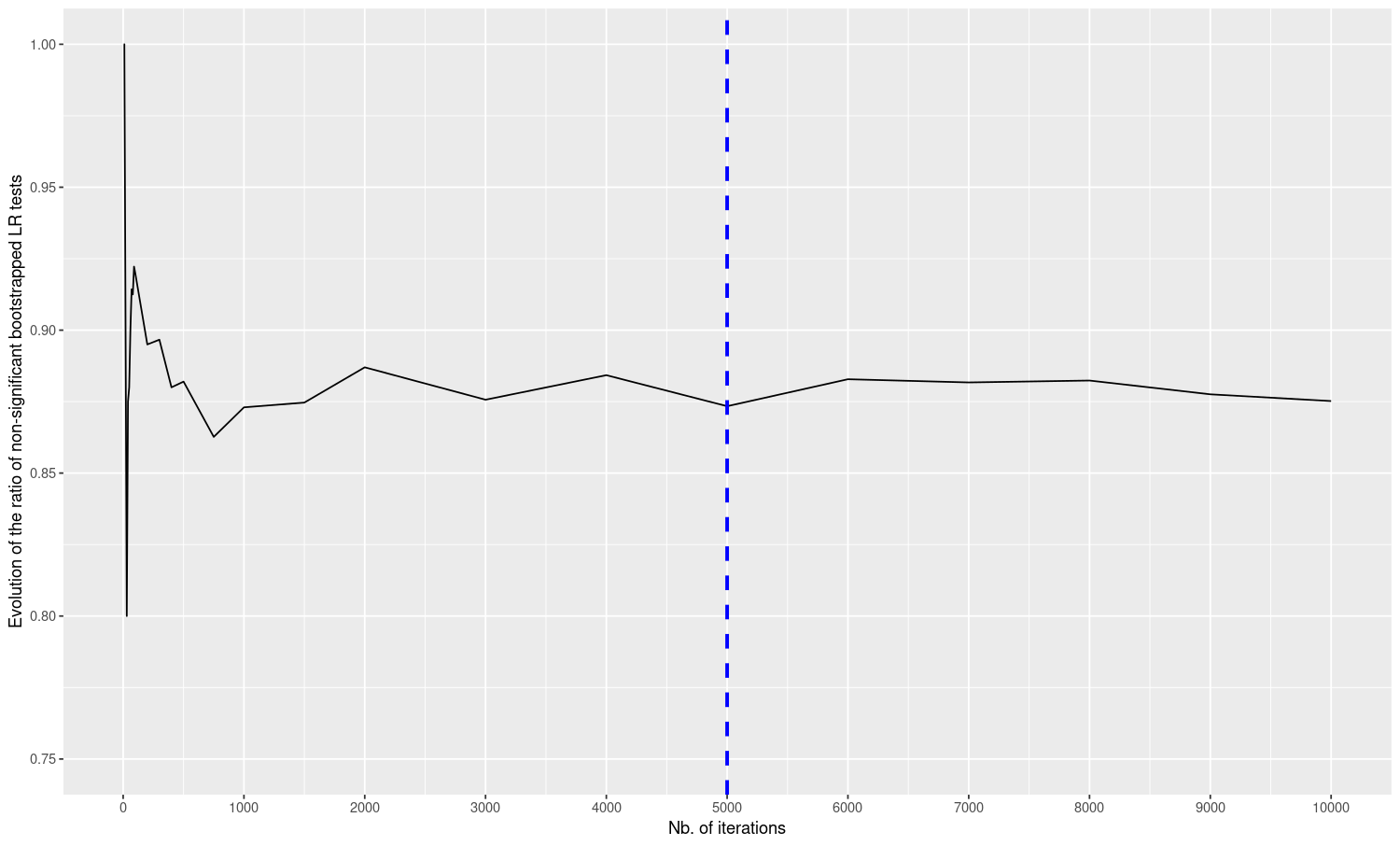
